# Taxonomy-aware, sequence similarity ranking reliably predicts phage-host relationships

**DOI:** 10.1101/2021.01.05.425417

**Authors:** Andrzej Zielezinski, Jakub Barylski, Wojciech M. Karlowski

**Affiliations:** Department of Computational Biology, Faculty of Biology, Adam Mickiewicz University Poznan, Uniwersytetu Poznanskiego 6, 61-614, Poznan, Poland; Molecular Virology Research Unit, Faculty of Biology, Adam Mickiewicz University Poznan, Uniwersytetu Poznanskiego 6, 61-614, Poznan, Poland

**Author notes:** To whom correspondence should be addressed. Andrzej Zielezinski, Wojciech M. Karlowski.

**Keywords:** phage-host prediction, phage, prokaryote, bacteria, virus, genome sequence, bioinformatics

## Abstract

**Background:** Similar regions in virus and host genomes provide strong evidence for phage-host interaction, and BLAST is one of the leading tools to predict prokaryotic hosts from phage sequences. However, BLAST-based host prediction has three major limitations: (i) top-scoring sequences do not always point to the actual host, (ii) mosaic phage genomes may match to many, typically related, bacteria, and (iii) phage and host sequences may diverge beyond the point where their relationship can be detected by a BLAST alignment.

**Results:** We created an extension to BLAST, named Phirbo, that improves host prediction quality beyond what is obtainable from standard BLAST searches. The tool harnesses information concerning sequence similarity and bacteria relatedness to predict phage-host interactions. Phirbo was evaluated on three benchmark sets of known phage-host pairs, and it improved precision and recall by 11-40 percentage points over currently available, state-of-the-art, alignment-based, alignment-free, and machine learning host prediction tools. Moreover, the discriminatory power of Phirbo for the recognition of phage-host relationships surpassed the results of other tools by at least 10 percentage points (Area Under the Curve = 0.95), yielding a mean host prediction accuracy of 57% and 68% at the genus and family levels respectively, and drops by 12 percentage points when using only a fraction of phage genome sequences (3 kb). Finally, we provide insights into a repertoire of protein and ncRNA genes that are shared between phages and hosts and may be prone to horizontal transfer during infection.

**Conclusions:** Our results suggest that Phirbo is a simple and effective tool for predicting phage host relationships.

## BACKGROUND

Prokaryotic viruses (phages) are the most abundant entities across all habitats and represent a vast reservoir of genetic diversity [1]. Phages mediate horizontal gene transfer and constitute a major selection pressure that shapes the evolution of bacteria [2]. Prokaryotic viruses also affect biogeochemical cycles and ecosystem dynamics by controlling microbial growth rates and releasing the contents of microbial cells into the environment [2, 3]. Moreover, phages play a key role in shaping the composition and function of the human microbiome in health and disease [4–6]. Recently, there has been renewed interest in phage therapy and phage-based biocontrol of harmful bacteria [7, 8] in medical treatment [9, 10] and the food industry [11, 12]. Hence, characterizing phage–host interactions is critical to understanding the factors that govern phage infection dynamics and their subsequent ecological consequences [13].

The scope of phage-host interactions is poorly understood, although it has been hypothesized that all prokaryotic organisms fall prey to viral attacks [1]. Methods for studying phage-host interactions primarily rely on cultured virus-host systems; however, recent *in silico* approaches suggest a much broader range of hosts may be susceptible to viral infections [14]. These methods predict prokaryotic hosts based on sequence composition [15, 16], direct sequence similarity between phages and hosts [14], analysis of CRISPR spacers or tRNAs [13, 17], as well as machine-learning approaches that integrate several sequence-based methods [18, 19].

Despite significant progress in phage-host predictions, the classic BLAST [20] algorithm is currently the most effective non-machine-learning method for identifying phage-host interactions [14, 15]. Depending on the dataset, the tool finds the correct genus level host for 40-60% of phages [14, 15]. The task of finding a host for a given phage using BLAST is conceptualized as obtaining the host sequence with the highest similarity to the query phage sequence. However, restricting host predictions to the first top-scored prokaryotic sequence has three limitations. First, the true host may not be the top-scoring match in the BLAST results. Second, selecting a prokaryotic host based on the first sequence assumes that a phage infects a single host. Although phages are generally host-specific, some may infect multiple host species [21, 22]. Finally, many distantly-related prokaryotic species may obtain a comparable BLAST score for a query phage due to spurious alignments. These ambiguous host predictions require further manual curation of the taxonomic or phylogenetic relationship between the top-scored prokaryotic species to select the true host(s).

We have addressed these issues by developing a simple extension to BLAST, named Phirbo, that exploits the information contained in the full BLAST results, rather than its top-ranking matches. Phirbo improved the accuracy of finding hosts, beyond what is found from the best BLAST match, by relating phage and host sequences through intermediate, common reference sequences that are potentially homologous to both phage and host queries. Subsequent quantification of the overlapping signals allows for the reliable prediction of phage-host interactions without the need for direct comparisons between the phage and host sequences and without any prior knowledge of their phylogenetic or taxonomic context.

## RESULTS

### Phirbo algorithm overview

Our algorithm is based on the assumption that the degree of similarity between phage and host sequences is proportional to the overlap between ranked similarity matches of each sequence to the same reference data set of prokaryotic sequences. Specifically, to compare a pair of phage (*P*) and host (*H*) sequences, we first perform two independent BLAST searches against the reference database of prokaryotic genomes (*D*)—one BLAST search for phage and the other for the host query (**Fig. 1a**). The two lists of BLAST results (**Fig. 1b**), *P* → *D* and *H* → *D*, contain prokaryotic genomes ordered by decreasing sequence similarity (i.e., bit-score). To avoid a taxonomic bias due to multiple genomes of the same prokaryote species, we rank prokaryotic species according to their first appearance in the BLAST list (**Fig. 1c**). In this way, both lists represent phage and host profiles consisting of the ranks of top-score prokaryotic species.

**Figure 1.**
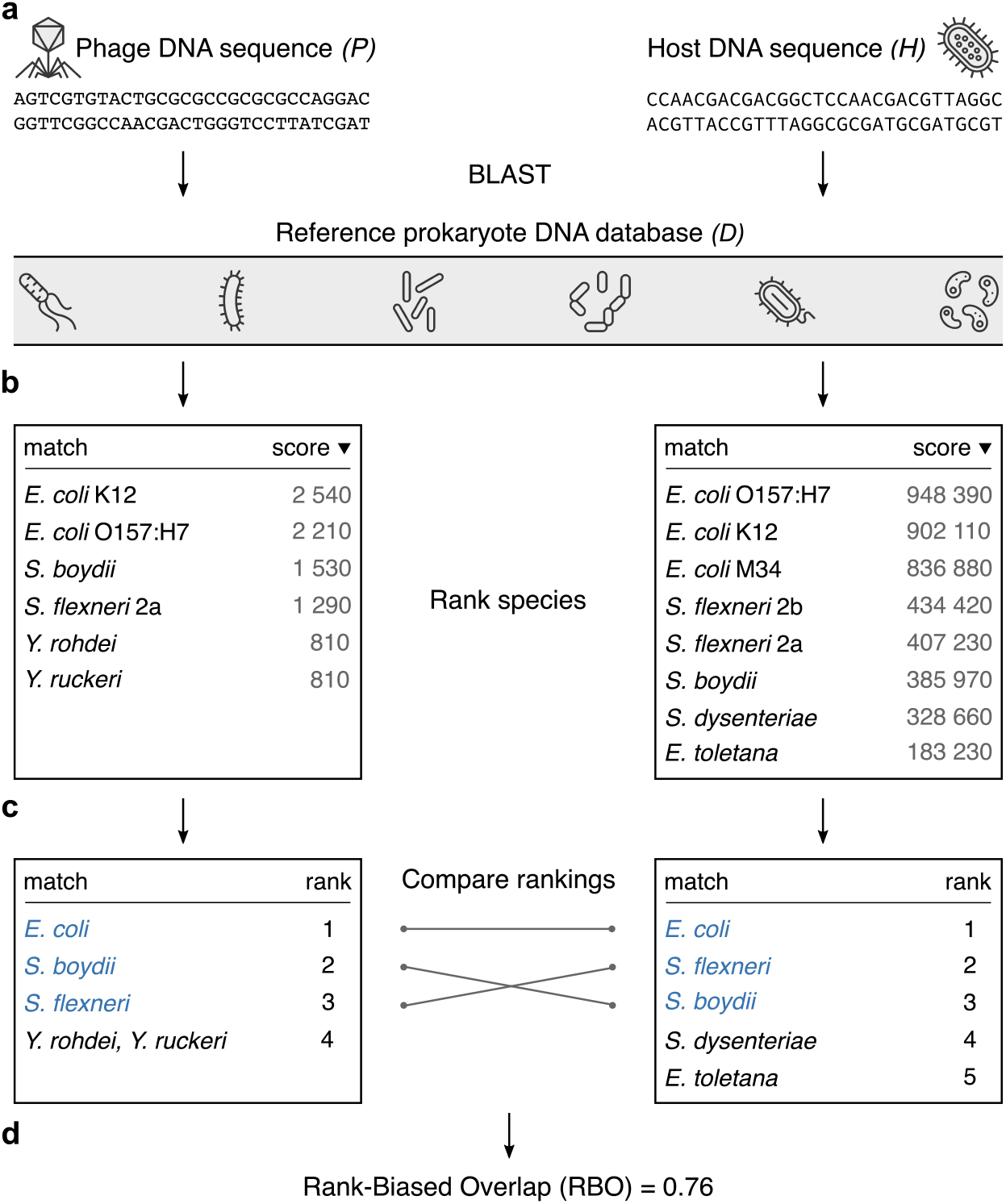
Calculation of the interaction score between phage and host sequences. **a.**The BLAST search of phage and prokaryote sequences against a reference dataset result in **b.** two BLAST lists containing prokaryote matches ordered by decreasing similarity (i.e., bit-score). **c.** BLAST lists were converted into rankings of prokaryote species. The ranked lists differ in content: *Yersinia rohdei* and *Y. ruckeri* are present in the first ranking list but absent in the second list, while *Shigella dysenteriae* and *Erwinia toletana* are only present in the second list. Two species, *Y. rohdei* and *Y. ruckeri*, from the first BLAST search have the same scores and are consequently tied for the same rank. **d.** An interaction score was calculated between two ranking lists using rank-biased overlap.

The properties of these lists (**Fig. 1c**) closely resemble the outcome of an Internet search and can be characterized by four features: (i) species listed at the top of each ranking are more important (similar) to the query than those listed at the bottom; (ii) the lists may not be conjoint (some species may appear in one ranking but not in the other); (iii) the ranking lists may vary in length (BLAST may return few prokaryotic matches in response to virus sequences in contrast to thousands of matches in cases of multiple-species prokaryotic families); (iv) two or more species from the database may achieve the same BLAST score and, therefore, occupy the same position on the ranking list (**Fig. 1c**). A recently introduced similarity measure used for comparing the rankings of Web search engine results [23], the Rank-Biased Overlap (RBO), satisfies these four conditions. The RBO algorithm starts by scoring the overlap between the sub-list containing the single top-ranked item of each list. It then proceeds by scoring the overlaps between sub-lists formed by the incremental addition of items further down the original lists. Each consecutive iteration has less impact on the final RBO score as it puts heavier weights on higher-ranking items by using geometric progression, which weighs the contribution of overlaps at lower ranks (see ‘Methods’). An overall RBO score falls between 0 and 1, where 0 signifies that the lists are disjoint (have no items in common) and 1 means the lists are identical in content and order. Our results indicate that the extent of the phage-host relationship can be estimated by the application of an RBO measurement to the ranking lists generated from BLAST results (**Fig. 1d**).

### Phirbo differentiates between interacting and non-interacting phage-host pairs

To assess the discriminatory power of Phirbo to recognize phage-host interactions, we used two published reference data sets: Edwards *et al*. (2016), which contains 2,699 complete bacterial genomes and 820 phages with reported hosts, and Galiez *et al.* (2017) that has 3,780 complete prokaryotic genomes and 1,420 phage genomes. For each data set, we compared the distribution of Phirbo scores between all known phage-host interaction pairs and the same number of randomly selected non-interacting phage-prokaryote pairs (**Additional file 1: Figure S1**). The scores obtained by Phirbo in both data sets separated the interacting from non-interacting phage-host pairs more than the BLAST scores. The median Phirbo score across interacting phage-host pairs was nearly 1,500 times greater than for non-interacting pairs, while the median BLAST score was three times higher for interacting pairs than non-interacting pairs (**Additional file 2: Table S1**). Both methods differentiated between interacting and non-interacting phage-host pairs with higher accuracy than WIsH — the state-of-the-art, alignment-free, host prediction tool [16].

To further examine the discriminatory power of Phirbo across all possible phage-prokaryote pairs, we used receiver operating characteristic (ROC) curves (**Fig. 2a,b**). The area under the ROC (AUC), which measured the discriminative ability between interacting and non-interacting phage-host pairs, was higher for Phirbo (AUC = 0.95) in the Edwards *et al*. and Galiez *et al*. data sets than for BLAST (AUC = 0.86) and WIsH (AUC = 0.78-0.79). An additional advantage of Phirbo was its capacity to score phage-host pairs whose sequence similarity could not be established by a direct BLAST comparison but, instead, through other, ‘intermediate’ prokaryotic sequences that were detectably similar to both phage and host query sequences. For example, BLAST did not provide scores for 20% of the interacting phage-host pairs in the Edwards *et al*. and Galiez *et al*. data sets due to alignment score thresholds (**Additional file 2: Table S2**). Using the same BLAST lists, Phirbo evaluated 99% of the interacting phage-hosts pairs. This high coverage indicated that nearly every pair of phage-prokaryote sequences could be related by at least one common prokaryotic sequence detectably similar to both the phage and host sequences.

**Figure 2.**
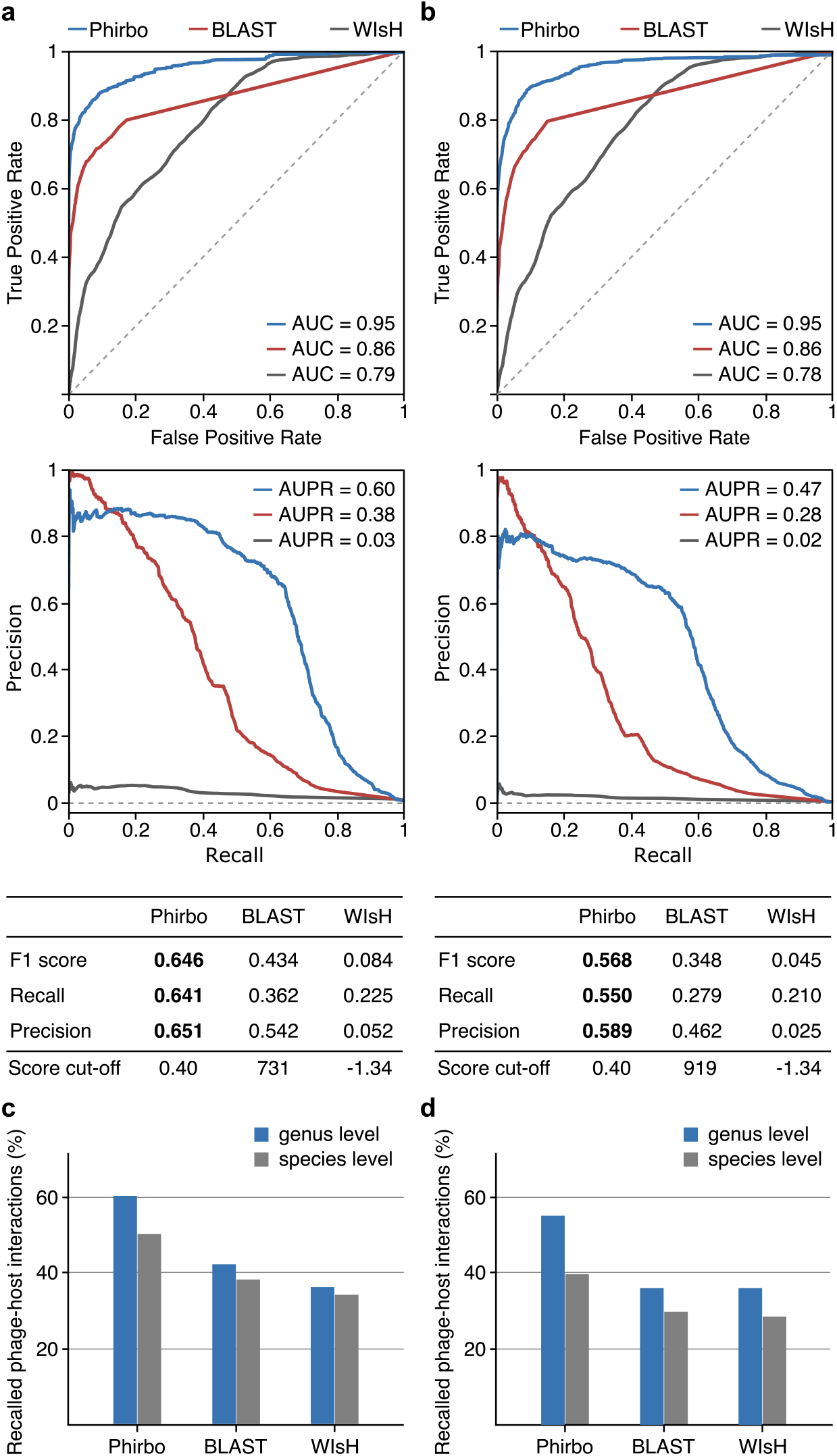
Host prediction performance of Phirbo, BLAST, and WIsH. The performance is provided by Receiver operating characteristic (ROC) and Precision-Recall (PR) curves and statistical measures (i.e., F1 score, precision, and recall) separately for **a.** Edwards *et al*. and **b.** Galiez *et al*. data sets. ROC curves and the corresponding area under the curve (AUC) display the classification accuracy of phage–host predictions across all possible phage-host pairs. Dashed lines represent the levels of discrimination expected by chance. Dashed lines in the PR-curve plots represent the levels of discrimination expected by chance. Score cut-offs for each tool were set to ensure the highest F1 score. **c.** and **d.** Number of correctly predicted phage-host interactions (%) in the Edwards *et al*. and Galiez *et al*. data sets, respectively. Bars indicate the number of phages for which a correct host was predicted at the species (blue bars) and genus (red bars) levels out of all phages in Edwards *et al*. (n = 820) and Galiez *et al*. (*n =*1,420).

### Phirbo has the highest host prediction performance

To evaluate host prediction performance, we used precision-recall (PR) curves, which provide more reliable information than ROC when benchmarking imbalanced data sets for which the non-interacting pairs vastly outnumber the interacting pairs [24, 25]. Accordingly, we plotted PR curves for Phirbo, BLAST, and WIsH predictions obtained from the Edwards *et al*. (**Fig. 2a**) and Galiez *et al*. (**Fig. 2b**) data sets. Overall, Phirbo performed better at host prediction at the species level than BLAST and WIsH, regardless of the data set. The area under the PR curve (AUPR), which summarized overall performance, was higher in Phirbo by 25 percentage points (AUPR = 0.56-0.65) than in BLAST (AUPR = 0.33-0.41).

Phirbo also reported the highest F1 score (an average of precision and recall [see ‘Methods’]) in the Edwards *et al*. and Galiez *et al*. data sets (**Fig. 2a,b**). Specifically, the precision and recall of Phirbo in predicting interacting phage-host pairs were 59-65% and 57-64%, respectively, while BLAST had precision and recall in the range of 28-43% (**Fig. 2a,b**). When setting a score cut-off that maximized the F1 score of each tool, Phirbo recalled 27-28% more interacting phage-host pairs than BLAST, and 34-41% more pairs than WIsH. Phirbo found the correct host at the species and genus levels for 38-50% and 55-60% of the analyzed phages, respectively (**Fig. 3c,d**). These results represent a 10% and 20% improvement over BLAST in the prediction of hosts at the species and genus levels, respectively.

**Figure 3.**
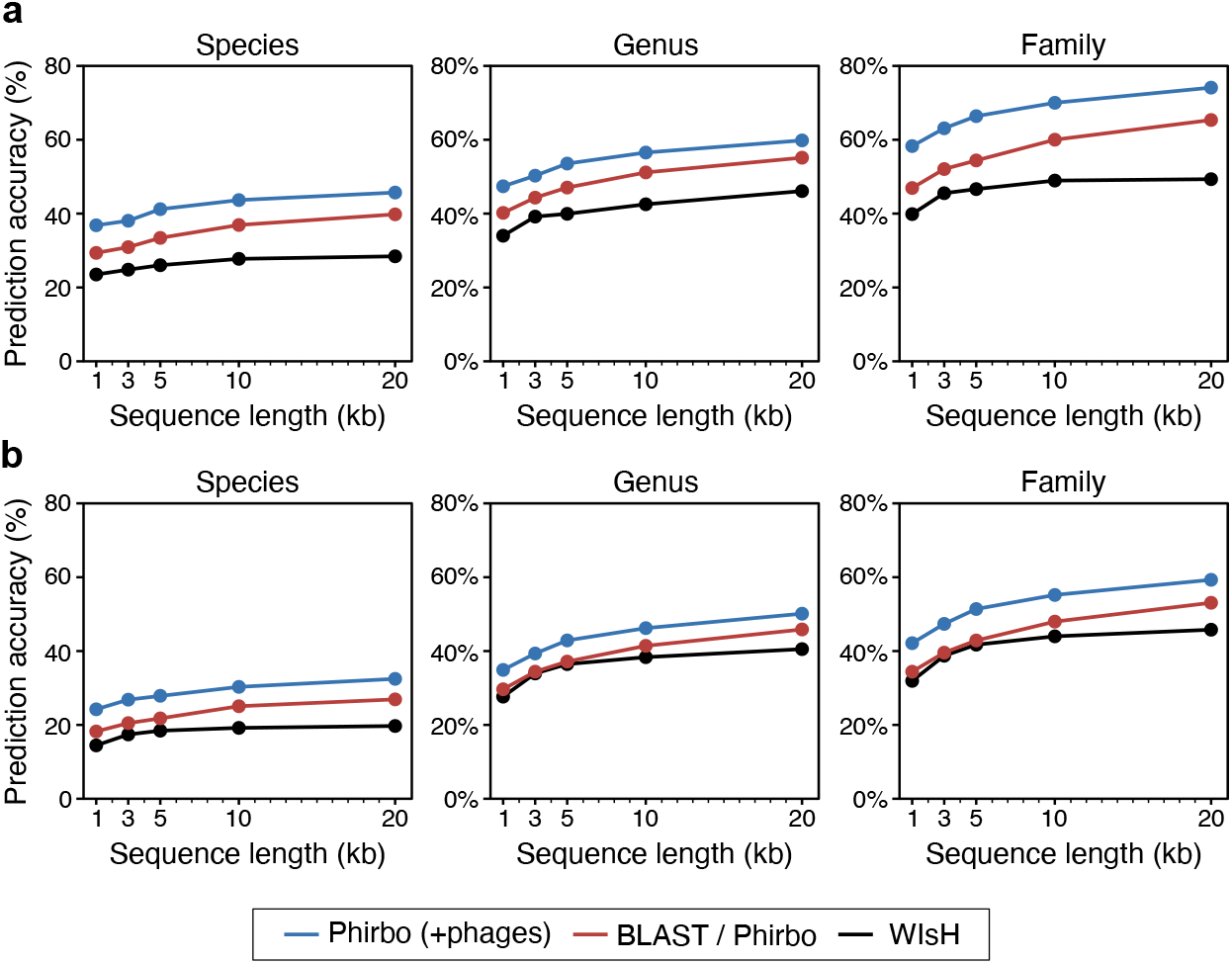
Host prediction accuracy over phage contig length. Prediction accuracy is provided separately for **a.** Edwards *et al*. and **b.** Galiez *et al*. data sets. Each complete virus genome was randomly subsampled 10 times for different sequence lengths (i.e., 20 kb, 10 kb, 5 kb, 3 kb, and 1 kb). Hosts were predicted on each subsampling replicate by selecting a prokaryotic sequence with the highest similarity to the query viral sequence. Points indicate the average of the resulting accuracies for all the viruses at a given subsampling length and host taxonomic level (i.e., species, genus, and family). An extended version of this figure containing host prediction accuracy values is provided in **Additional file2: Table S4**.

### Phirbo preserves BLAST top-ranked host predictions

We further evaluated the host prediction accuracy of Phirbo by selecting a top-scored prokaryotic sequence for each phage without thresholds on score values [14–16, 18]. Briefly, host prediction accuracy is calculated as the percentage of phages whose predicted hosts have the same taxonomic affiliation as their respective known hosts (if multiple top-scoring hosts are present, the prediction is scored as correct if the true host is among the predicted hosts). Phirbo restored all hosts predicted by BLAST in the datasets by Edwards *et al.* and Galiez *et al*., achieving the same prediction accuracy as BLAST across all taxonomic levels (**Table 1**). Of note, BLAST found multiple different host species with equal scores for 14 phage genomes. This was observed in phages infecting bacteria from the Enterobacteriaceae family and the Rhodococcus and Bacillus genera. However, Phirbo assigned the highest score to the correct host species (**Additional file 2: Table S3**). Additionally, it refined the host prediction for the Cronobacter phage ENT39118 sequence, which BLAST assigned to the *Escherichia coli* genome. Phirbo revealed *Cronobacter sakazaki* as the primary host species, as the BLAST list of the Cronobacter phage is more similar in content and order to the BLAST list of *C. sakazaki* (Phirbo score = 0.50) than *E. coli* (Phirbo score: 0.48) (**Additional file 1: Figure S2**).

**Table 1.** Host prediction accuracies (%) for phage and host genomes from the data sets by Edwards *et al*. [14] and Galiez *et al*. [16].

| Data set | Method | Species | Genus | Family | Order | Class | Phylum |
| --- | --- | --- | --- | --- | --- | --- | --- |
| Edwards *et al.* (2016) | WIsH | 28 | 44 | 50 | 53 | 62 | 70 |
|  | BLAST | 43 | 59 | 71 | 78 | 87 | 96 |
|  | Phirbo* | 43 | 59 | 71 | 78 | 87 | 96 |
|  | Phirbo (+phages)<sup>†</sup> | **48** | **63** | **75** | **82** | **90** | **97** |
| Galiez *et al.* (2017) | WIsH | 21 | 44 | 48 | 53 | 68 | 77 |
|  | BLAST | 31 | 53 | 62 | 68 | 88 | 95 |
|  | Phirbo* | 31 | 53 | 62 | 68 | 88 | 95 |
|  | Phirbo (+phages)<sup>†</sup> | **35** | **56** | **65** | **72** | **90** | **96** |

The highest accuracies among the methods for each taxonomic level are in bold.
\* Phirbo scores were calculated using rank-biased overlap (RBO) between BLAST lists containing prokaryotic sequences. Specifically, the BLAST database contained 2,699 sequences of bacterial genomes in the Edwards *et al.* data set, and 3,780 sequences of bacterial and archaeal genomes in the Galiez *et al.* data set.
<sup>†</sup> Phirbo scores were calculated using RBO between BLAST lists containing both prokaryotic and phage sequences.

As Phirbo links phage to host through common sequences, the content of the sequence database was the main factor defining host prediction quality. Since the similarity between viruses may indicate a common host [18, 26], we expanded the two BLAST databases of prokaryotic sequences obtained from Edwards *et al*. and Galiez *et al*. by phage sequences (*n* = 820 and *n* = 1420, respectively), and recalculated Phirbo scores between every phage-prokaryote pair. The phage-host linkage through homologous prokaryotic and phage sequences increased the host prediction accuracy of Phirbo at all taxonomic levels, allowing correct identification of hosts at the genus level for 56-63% of phages (**Table 1**). Specifically, Phirbo refined BLAST mis-predictions for 55 phage genomes and showed which sequences demonstrated low similarity to the sequences of their host species. The direct BLAST alignments of these phage sequences, and the sequences of their corresponding hosts, obtained significantly lower scores than alignments obtained by the other known phage-host pairs (*P* = 1.9 × 10^−45^, Mann–Whitney U test). Notably, Phirbo also assigned correct host species for 18 phages whose hosts were not reported in the BLAST results, mainly Chlamydia species, *Vibrio cholerae*, and the opportunistic pathogen, *Acinetobacter baumannii*.

### Phirbo is suitable for incomplete phage sequences

We tested the robustness of our host prediction algorithm to fragmentation of the phage sequence. Following earlier studies [15, 16, 18], phage genomes from Edwards *et al*. and Galiez *et al*. data sets were randomly subsampled to generate contigs of different lengths (20 kb, 10 kb, 5 kb, 3 kb, and 1 kb) with 10 replicates. Host prediction accuracy was calculated as the mean percentage of phages whose predicted hosts had the same taxonomic affiliation as their respective known hosts (**Fig. 3**). Although Phirbo achieved equal host prediction accuracy with BLAST across all contig lengths, it had substantially higher overall performance in terms of AUC and AUPR (**Fig. S3**; *P*□<□10^−5^, Wilcoxon signed-rank test). Surprisingly, BLAST-based methods obtained higher host prediction accuracy across all contig lengths compared to WIsH, a tool designed to predict the hosts of short viral contigs (**Fig. 3**).

The host prediction accuracy of Phirbo was examined using the expanded BLAST database of both prokaryotic and phage full-length sequences. To ensure fairness, for each tested phage contig we removed its corresponding full-length sequence from the BLAST database and recalculated Phirbo scores between the phage contig and every prokaryotic sequence. This approach outperformed BLAST at every contig length across all taxonomic levels in both data sets (**Fig. 3**). Generally, the host prediction accuracy of Phirbo improved by 5-11 percentage points compared to the BLAST results. For example, when the contig length was 3 kb, the prediction accuracy of Phirbo was 8-11% higher than BLAST at the family level, and 8-17% higher than WIsH (**Fig. 3**; **Additional file 2: Table S4**). Phirbo also achieved the highest AUC and AUPR scores when discriminating between interacting and non-interacting phage-host pairs (**Additional file 1: Figure S3**).

### Phirbo uses multiple protein and non-coding RNA signals for host prediction

We investigated the sequence information used by BLAST and Phirbo for host prediction. For each phage that was correctly assigned to the host species by both tools (*n* = 485), we calculated the fraction of the phage genome that was included in the segments aligned with prokaryotic sequences (sequence coverage). This analysis revealed that our tool used three times more phage sequence (median sequence coverage: 35%) than BLAST (12%) (**Additional file 1: Figure S4**; *P* < 10^−15^, Wilcoxon signed-rank test). This increased sequence coverage indicates that different genome regions of the phages map to the genomes of prokaryotic species other than the host species. For 214 of the 485 phages, more than half of their genomes were aligned to genomes of their host species (**Additional file 2: Table S5**). Such large regions of homology are likely prophages or phage debris left by large-scale recombination events during phage replication. The observed high sequence coverage points to the virus taxa, known for their temperate lifestyle and frequent recombination with host genomes (i.e., Siphoviridae family as well as the Peduovirinae and Sepvirinae subfamilies).

To further examine the properties of sequences that may be exchanged between a phage and its host, we selected a population of phages with sequence coverage below 50% (*n* = 271). These phages, which are less likely to represent complete prophages, belong to 16 viral families (**Additional file 2: Table S6**). Next, we re-annotated the genomic sequences of the phages to find putative protein and non-coding RNA (ncRNA) genes. Phage sequence regions used by Phirbo for host predictions were significantly enriched (*P* < 10^−5^) in more than a hundred protein families of known or probable function. In contrast, only half of the protein families were used in BLAST-based host predictions (**Additional file 2: Table S7**). The protein families used by Phirbo covered most of the processes of the viral life cycle including DNA replication, cell lysis, recombination, and packaging of the phage genome (**Fig. 4**). In contrast to BLAST, Phirbo also exploited the information contained in phage ncRNAs while assigning phages to host genomes. The vast majority of these ncRNAs (>90%) were tRNAs, which showed significant overrepresentation in the phage sequence fragments used by Phirbo (*P* = 6 × 10^−12^) (**Additional file 2: Table S8**). The remaining ncRNAs belonged to group I introns (3%), RNAs associated with genes associated with twister and hammerhead ribozymes (1%), skipping-rope RNA motifs (1%), and 12 less abundant RNA families.

**Figure 4.**
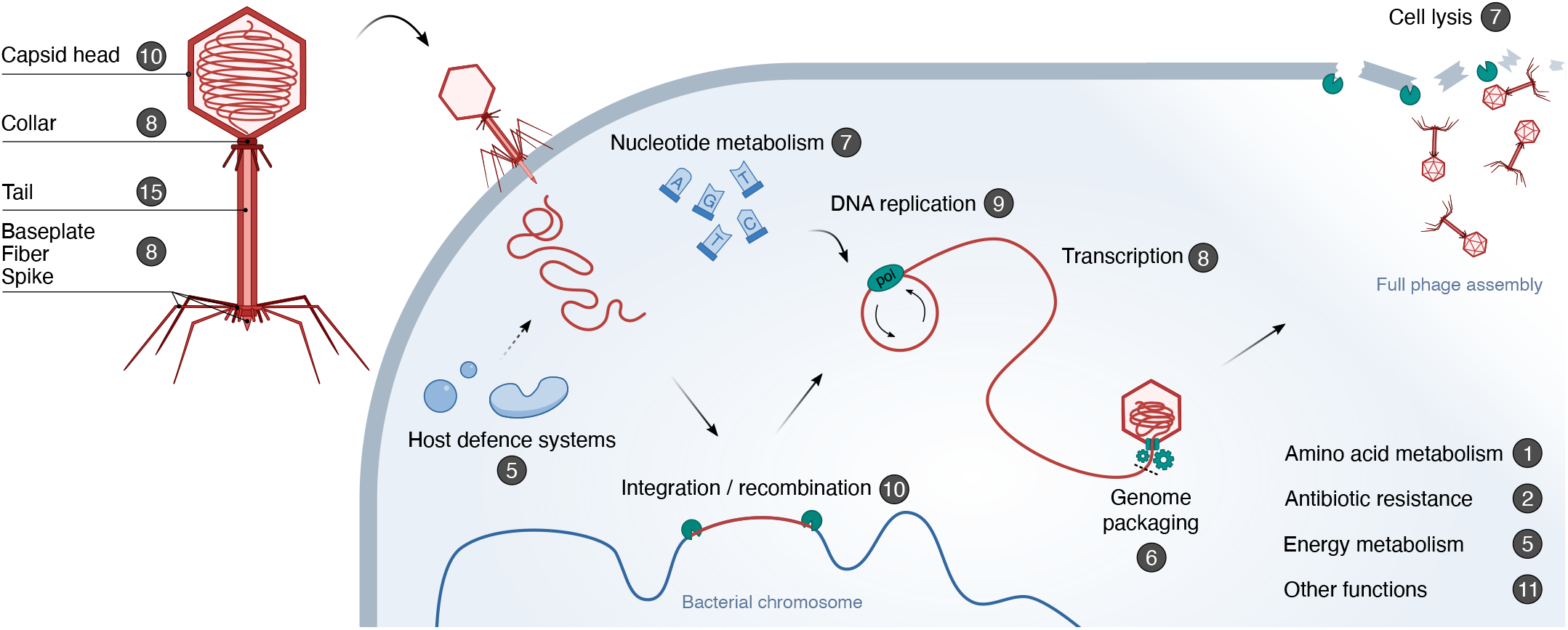
Functional classification of phage coding sequences used by Phirbo for host prediction. Protein families (pVOGs) were classified into 15 functions (e.g., DNA replication, transcription). Numbers in the dark circles indicate the number of different pVOGs related to a given function. An extended version of this figure containing the list of pVOGs is provided in **Additional file 2: Table S7**.

### Phirbo has higher precision and recall than VirHostMatcher-Net and PHP

We tested Phirbo against two machine-learning host prediction tools, VirHostMatcher-Net [18] and Prokaryotic virus Host Predictor (PHP) [19]. VirHostMatcher-Net predicts phage–host interactions using multiple virus-host and virus-virus sequence similarity features including BLAST. PHP utilizes a Gaussian model based on differences of *k*-mer frequencies between viral and host genomic sequences. We benchmarked Phirbo, VirHostMatcher-Net, and PHP using the Wang *et al.* (2020) data set of 1,462 phages (*W*) and 62,493 candidate hosts. Analogously, we calculated host prediction accuracy for each tool by selecting a top-scored prokaryotic sequence for each phage (**Table 2**). Although Phirbo outperformed PHP across all taxonomic levels (from species to phylum), it had lower prediction accuracy than VirHostMatcher-Net at taxonomic levels from up to the class level.

**Table 2.** Host prediction accuracies (%) for phage and host genomes from the Wang *et al.* [18] data set.

| Data set | Method | Species | Genus | Family | Order | Class | Phylum |
| --- | --- | --- | --- | --- | --- | --- | --- |
| $W$ (1462 phages) | Phirbo | 32 | 52 | 64 | 69 | 78 | **87** |
|  | VirHostMatcher-Net | **44** | **59** | **70** | **78** | **84** | 86 |
|  | PHP | 20 | 44 | 62 | 67 | 77 | 85 |
| $W_{species}$ (451 phages) | Phirbo | **32** | **62** | **71** | **80** | **87** | **92** |
|  | VirHostMatcher-Net | 11 | 51 | 58 | 71 | 82 | 85 |
|  | PHP | 12 | 34 | 49 | 68 | 82 | 91 |
| $W_{genus}$ (261 phages) | Phirbo | **37** | **56** | **69** | **77** | **84** | **89** |
|  | VirHostMatcher-Net | 20 | 34 | 48 | 64 | 78 | 81 |
|  | PHP | 14 | 25 | 44 | 67 | 77 | 87 |
| $W_{family}$ (171 phages) | Phirbo | **34** | **56** | **66** | **74** | **83** | **89** |
|  | VirHostMatcher-Net | 22 | 39 | 42 | 62 | 79 | 80 |
|  | PHP | 10 | 26 | 37 | 66 | 79 | 88 |

The highest accuracies among the methods for each data set and taxonomic level are in bold.

Although VirHostMatcher-Net and PHP were trained and tested on mutually exclusive sets of phages [18, 19], both data sets contained phages that have high sequence similarity and infect the same host species. To minimize the effect on the benchmark results of these potentially crossmatching sequences, we performed a more stringent test that gradually separated the testing phage sequences from the training data set. Specifically, we assembled three subsets (*W_species_*, *W_genus_*, and *W_family_*) from the Wang *et al*. virus set (1,462 phages). *W_species_* consisted of all phages for which host specificity at the species level was different than for viruses in the original training set. Correspondingly, *W_genus_*, and *W_family_* sets had different host genera or families, respectively.

Across the three data sets, Phirbo achieved the highest host prediction accuracy at all taxonomic levels; for example, it recalled the correct host genus for 56-62% of phages—outperforming VirHostMatcher-Net (*n* = 34-51%) and PHP (*n* = 25-34%) (**Table 2**). Phirbo was also markedly robust in regard to data set heterogeneity, as predictions across all four data sets varied significantly less, particularly at the species, genus, and family levels (standard deviation = 2-4%), than results of VirHostMatcher-Net (11-14%) and PHP (4-11%).

To compare the performance of Phirbo, VirHostMatcher-Net, and PHP at different score thresholds, we plotted ROC and PR curves for all four data sets: *W*, *W_species_*, *W_genus_*, and *W_family_* (**Fig. 5**). Phirbo discriminated between interacting and non-interacting phage-host pairs with higher accuracy (AUC = 0.95) than VirHostMatcher-Net (AUC = 0.86-0.9) and PHP (AUC = 0.83-0.88) (**Fig. 5a**). Our tool also provided a better precision-recall trade-off (AUPR = 0.31-0.45) than VirHostMatcher-Net (AUPR = 0.07-0.34) and PHP (0.02-0.04) (**Fig. 5b**).

**Figure 5.**
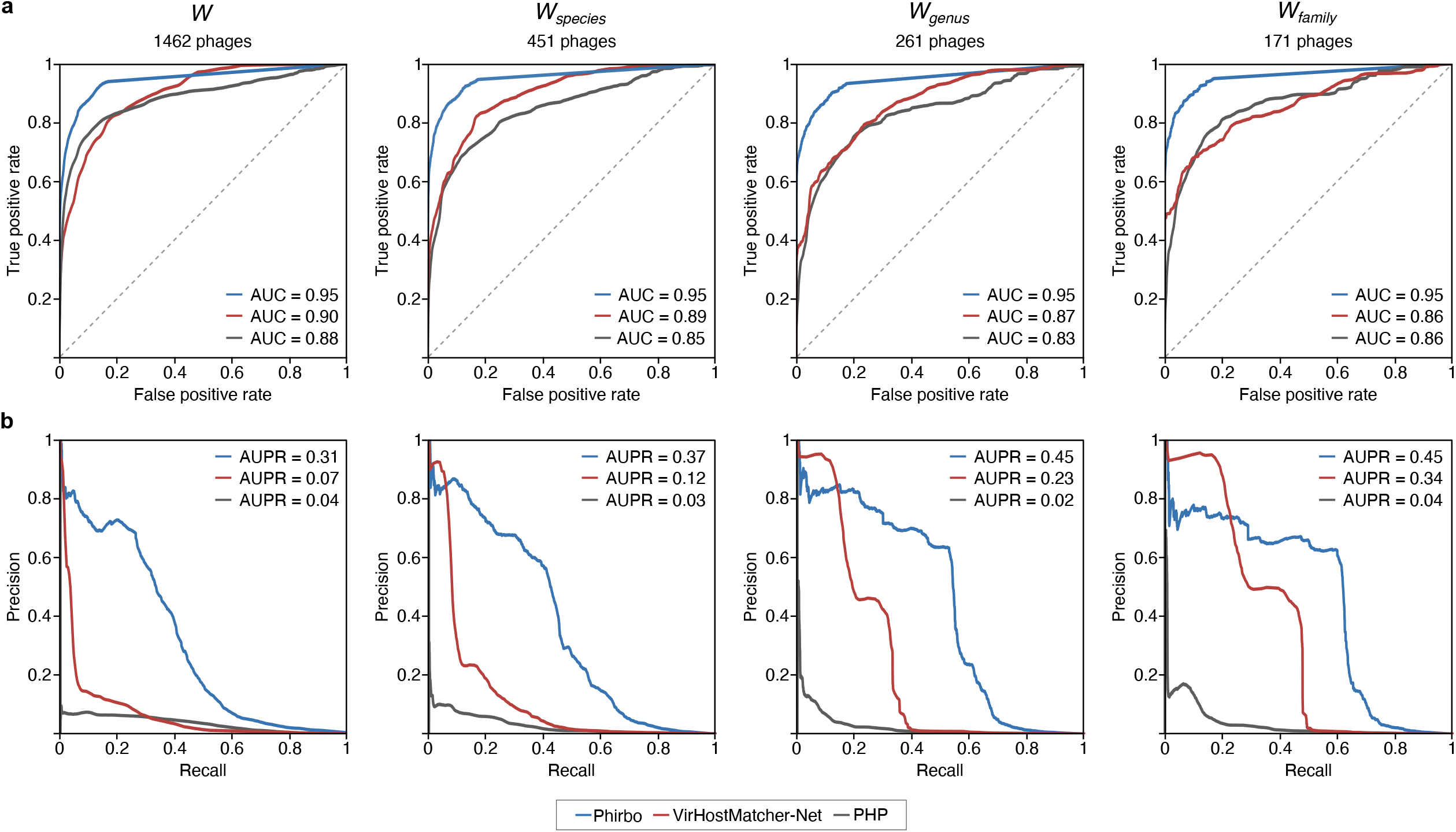
Host prediction performance of Phirbo, VirHostMatcher-Net, and PHP on the Wang *et al*. data set. The performance was evaluated using **a.** Precision-Recall (PR) curves and **b.** Receiver operating characteristic (ROC) curves.

### Implementation and availability

Predicting hosts from phage sequences using BLAST is accomplished by querying phage sequences against a database of candidate hosts. However, Phirbo also uses information about sequence relatedness among prokaryotic genomes. Therefore, it requires ranked lists of prokaryote species generated by BLAST for the phage and host genomes. The computational cost of querying every host sequence against the database of all candidate hosts using BLAST may still be a limiting factor. However, for mass host searches, the computational cost of all-versus-all host comparisons becomes marginal, as it must be done only once. After the relatedness among host genomes is established, the time required for Phirbo host predictions is negligibly higher than the time for typical BLAST-based host predictions. For example, running Phirbo between ranked lists of host species for 1,462 phages and 62,493 candidate hosts from Wang *et al*. (resulting in ∼91 million phage-host comparisons) took 2 hours on a 16-core 2.60GHz Intel Xeon.

As Phirbo operates on rankings, BLAST can be replaced by an alternative sequence similarity search tool to reduce the time to estimate homologous relationships between host genomes. For instance, Mash [27] computed host relationships in 2 hours for the Wang *et al*. data set that encompassed 62,493 bacterial and archaeal genomes (see ‘Methods’). The host prediction performance of Phirbo using BLAST-based rankings for phages and Mash-based rankings for host genomes is comparably high to the performance of Phirbo predictions using BLAST rankings for both phage and host genomes (**Additional file 2: Table S10**).

We envisage Phirbo as a natural extension to standard BLAST-based host predictions. The Phirbo tool is written in Python and freely available at https://github.com/aziele/phirbo/.

## DISCUSSION

The identification of similar sequence regions between host and phage genomes using BLAST has been a baseline for the identification of putative virus-host connections in numerous metagenomic projects [13, 28, 29]. However, a BLAST search requires regions with significant similarity between the query phage and host [14–16]. Yet, many phage and host sequences lack sufficient similarity and escape detection with standard BLAST searches. To tackle this issue, alignment-free tools have been developed to predict hosts from phage sequences [14–16, 30]. The rationale behind these tools is based on the observation that viruses tend to share similar patterns in codon usage or short sequence fragments with their hosts [14–16]. As virus replication is dependent on the translational machinery of its host, some phages adapt their codon usage to match the availability of tRNAs during viral replication in the host cell [31–33]. Similar oligonucleotide frequency use may be driven by evolutionary pressure on the virus to avoid recognition by host restriction enzymes and CRISPR/Cas defense systems [32, 34]. Although state-of-the-art alignment-free tools (i.e., WIsH [16] and VirusHostMatcher [15]) can rapidly assess sequence similarity between any pair of phage and prokaryote sequences, they are less accurate for host prediction than BLAST [14, 15]. The relatively high accuracy of BLAST suggests that localized similarities of genetic material may be a stronger indication of phage-host interactions than global convergence of their genomic composition. This evidence comes in the form of protein-coding DNA fragments and non-coding RNAs. The latter group is dominated by tRNA genes, which are strongly over-represented in direct BLAST alignments between phages and their hosts, and are even more prevalent among indirect connections used by Phirbo. This may be important, as previous studies have shown that not all phage tRNA genes come directly from their hosts. Some appear to be derived from genomes of other, often distantly related, bacteria and may be the result of earlier evolutionary events [35]. For protein-coding genes, a more diverse picture emerges. Proteins rich in phage-host BLAST alignments can be assigned into different functional categories including phage virion components, replication-related proteins, regulatory factors, and proteins involved in the metabolism of the host. The transfer of some over-represented families in phages and/or prophages has been previously reported (e.g., lytic proteins, DNA replication and recombination proteins, and enzymes involved in nucleotide and energy metabolisms [36]) and some of these genes are connected with the phage-host range [37, 38]. However, no clear pattern emerges after analyzing the functions of the remaining, over-represented proteins.

In this study, we attempted to expand the information content of a single local alignment of phage and host sequences by incorporating the results of multiple alignments between a phage sequence and different prokaryotic genomes. This approach may more closely resemble a manual assignment of phage-host pairs, where an expert analyst not only considers a top-ranked matching prokaryote in the BLAST results, but also uses the information contained in other, less significant, matches and their sequence and taxonomic similarity. Through a taxonomically-aware stratification scheme, this approach tracks the multilateral dynamics of horizontal gene transfer. Therefore, we propose to relate phage and host sequences through multiple intermediate sequences that are detectably similar to both the phage and host sequences. By linking phage and host sequences through similar sequences, Phirbo achieved a more comprehensive list of phage-host interactions than BLAST. Simultaneously, Phirbo was capable of assessing almost all phage-host pairs, bringing the method closer to alignment-free tools, which compute scores between all possible phage and host pairs. Thus, our approach can be directly applied to different phage and prokaryote data sets without training or optimizing the underlying RBO algorithm.

## CONCLUSIONS

Our results show that expanding the information obtained from plain similarity comparisons by incorporating taxonomically-grounded measurements of phage-host similarity leads to improved precision and recall of phage-host predictions. The Phirbo method provides the phage research community with an easy-to-use tool for predicting the host genus and species of query phages, which is usable when searching for phages with appropriate host specificity and for correlating phages and hosts in ecological and metagenomic studies.

## METHODS

### Virus and prokaryotic host data sets

The data sets analyzed in this study were retrieved from three previously published phage-host studies [14, 16, 18]. The first set [14] contained 2,699 complete bacterial genomes obtained from NCBI RefSeq and 820 RefSeq genomes of phages for which the host was reported. The data set encompassed 16,757 known virus-host interaction pairs and 2,196,424 pairs for which interaction was not reported (non-interacting phage-host pairs). The second data set [16] contained 3,780 complete prokaryotic genomes of the KEGG database and 1420 phages for which host species were reported in the RefSeq Virus database. The data set consisted of 26,024 interacting- and 5,341,576 non-interacting virus-host pairs. The third data set [18] included 1,462 phages and 62,493 candidate hosts encompassing 113,250 interacting- and 91,251,516 non-interacting virus-host pairs.

### Phirbo score

The interaction score for a given phage-host pair was calculated using the Ranked-Biased Overlap (RBO) measure. RBO [23] is a measurement of rank similarity that compares two lists of different lengths (giving more attention to high ranks on the lists). RBO ranges from 0 to 1, where a greater value indicates greater similarity between lists. Equation 1 was used for the calculation of the RBO value between two ranking lists, *S* and *T*.

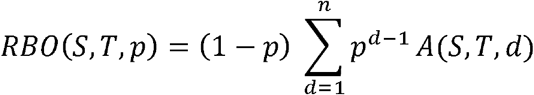

where the parameter *p* (0 < *p* < 1) determines how steeply the weight declines (the smaller the *p*, the more top results are weighted). When *p* = 0, only the top-ranked item is considered, and the RBO score is either zero or one. In this study, we set *p* to 0.75, which assigned ∼98% of the weight to the first 10 hosts. *A(S, T, d)* is the value of overlap between the two ranking lists, *S* and *T*, up to rank *d*, calculated by Eq. 2. *n* is the number of distinct ranks on the ranking list.

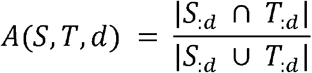

where *S_:d_* and *T_:d_* represents the elements present in the first *d* ranks of lists *S* and *T*, respectively.

### Host prediction tools

The host prediction tools BLAST [20], WIsH [16], and Phirbo were run separately in the Edwards *et al*. and Galiez *et al*. data sets. For each tool, sequence similarity scores were calculated across all combinations of phage-host pairs. BLAST 2.7.1+ [39] was run with default parameters (task: blastn, *e*-value threshold = 10) to query each phage sequence against a database of candidate host genomes. For each BLAST alignment, the highest bit-score between every phage-host pair was reported (for phage-host pairs that were absent in the BLAST results, a bit-score of 0 was assigned). For RBO host prediction, an additional BLAST search was performed to establish ranked lists of genetically similar host genomes. Specifically, a nucleotide BLAST was run with default parameters to query each host sequence against a database of candidate host genomes. As an alternative to BLAST, Mash 2.1 [27] was used with default parameters (*k*-mer size = 21, sketch size = 1,000) to establish ranked lists for each host by comparing its sequence against the database of candidate host genomes. RBO scores were calculated between all pairwise combinations of phage and host ranking lists. WIsH 1.0 [16] was used with default parameters to calculate log-likelihood scores between all pairwise combinations of phage-host sequences. VirHostMatcher-Net 1.0 [18] and Prokaryotic virus Host Predictor (PHP) [19] were run using default parameters.

### Evaluation metrics

The metrics of host prediction performance were calculated using sklearn (i.e., AUC, AUPR, recall, precision, specificity, and accuracy) [40]. Optimal score thresholds to calculate recall, precision, specificity, and accuracy were computed as maximizing the F1 score, an accuracy metric, which is the harmonic mean of precision and recall. Host prediction accuracy was evaluated analogous to previous studies [14, 16, 18]. Specifically, for each query phage, the host with the highest score to the query virus was selected as the predicted host. In cases where multiple hosts were predicted, the prediction was scored as correct if the correct host was among the predictions. The prediction accuracy was calculated at each taxonomic level as the percentage of viruses whose predicted hosts shared a taxonomic affiliation with known hosts.

### Phage genome annotation

To define phage genes potentially exchanged between phage and host genomes, we re-annotated 485 phage genomes that were correctly assigned to host species by both Phirbo and BLAST. The genes were classified into predefined pVOGs (prokaryotic Virus Orthologous Groups) [41] and RNA families [42]. Briefly, open reading frames (ORFs) in the analyzed 485 phage genomes were identified using Transeq from EMBOSS [43]. The ORFs were then assigned to the respective orthologue group by HMMsearch (*e*-value < 10^−5^) against the database of Hidden Markov Models (HMMs) created for every of 9,518 pVOG alignments using HMMbuild of HMMER v3.3.1 [44]. Non-coding RNAs (ncRNAs) were predicted in the phage genomes (*e*-value < 10^−5^) using Rfam covariance models v14.3 [42] and the Infernal tool v1.1.3 [45]. We counted the number of times each pVOG and Rfam term was present in phage sequences used by BLAST and Phirbo during host prediction. To determine whether the observed level of pVOG/Rfam counts was significant within the context of all the terms within the phage genome, we calculated the *p*-value using the hypergeometric distribution implemented in Scipy [46].

## LIST OF ABBREVIATIONS

AUC: Area under the ROC curve
AUPR: Area under the precision-recall curve
PHP: Prokaryotic virus Host Predictor
RBO: Ranked-biased overlap
ROC: Receiver operating characteristic

## ACKNOWLEDGMENTS

We thank Bas Dutilh, Rob Edwards, Clovis Galiez, Johannes Söding, Fengzhu Sun, and Weili Wang for providing us with the benchmark data sets used in their studies. We likewise acknowledge William Webber for assistance with modifying the RBO formula to account for tied ranks. The computations were performed at the Poznan Supercomputing and Networking Center.

## AUTHORS’ CONTRIBUTIONS

AZ conceived the project and designed the experiments. AZ wrote Phirbo and tested its performance. WMK provided the conceptual framework for sequence comparisons through intermediate sequences and reviewed the software and manuscript. AZ and JB analyzed the results and wrote the paper. All authors read and approved the final manuscript.

## AVAILABILITY OF DATA AND MATERIALS

Phirbo is available at https://github.com/aziele/phirbo.

## ETHICS APPROVAL AND CONSENT TO PARTICIPATE

Not applicable

## CONSENT FOR PUBLICATION

Not applicable

## ADDITIONAL FILES

### Additional file 1

**Figure S1.** Discriminatory power of Phirbo, BLAST, and WIsH scores to differentiate between interacting and non-interacting phage-host pairs. Phage-host pairs were obtained from **a.** Edwards *et al*. and **b.** Galiez *et al*. data sets. Box plots show the distribution of scores for all interacting phage-host pairs (*n* = 16,757 and *n* = 26,024 in Edwards *et al.* and Galiez *et al.*, respectively) and the same number of randomly selected, non-interacting phage-host pairs. The horizontal line in each box displays the median; boxes display the first and third quartiles; whiskers depict lowest and highest non-outlier scores (details of distributions including outliers are provided in **Additional file 2: Table S1**).

**Figure S2.** Host predictions for Cronobacter phage ENT39118 (RefSeq accession: NC_019934) using **a.** BLAST and **b.** Phirbo. Querying the Cronobacter phage sequence with a BLAST search against the host database returned the genomic sequence of *Escherichia coli* (NC_017641) as the best match (bit-score = 14,588), and *Cronobacter sakazakii* (NC_009778) as the second-best match (bit-score = 14,020). Phirbo predicted *Cronobacter sakazakii* as the top-score host for the Cronobacter phage due to the highest extent of overlap between the top-ranking BLAST matches of each sequence (NC_019934 and NC_009778) of the same database. For clarity, only the first ten BLAST matches are shown.

**Figure S3.** Host prediction performance of Phirbo, BLAST and WIsH over phage contig length in terms of **a.** Area under the curve (AUC) and **b.** Area under the precision-recall curve (AUPR). Bars indicate the AUC or AUPR averaged across 10 replicates at a given subsampling length of phage sequence.

**Figure S4.** Scatter plot of the phage sequence coverage used in host predictions of Phirbo versus that of BLAST. Each dot represents a phage genome.

### Additional file 2

**Table S1.** Distribution of Phirbo, BLAST and WIsH scores among interacting and non-interacting phage-host pairs obtained from Edwards *et al*. and Galiez *et al*. data sets. Score ranges were summarized separately for 16,757 interacting and non-interacting phage-host pairs from Edwards *et al*., and 26,024 interacting and non-interacting phage-host pairs from Galiez *et al*.

**Table S2.** Number of phage-host pairs evaluated by Phirbo, BLAST, and WIsH in Edwards *et al*. and Galiez *et al*. data sets.

**Table S3.** Phages assigned by BLAST to multiple, equally-scored host species. Phirbo differentiated between host species and provided the highest score to primary host species.

**Table S4.** Host prediction accuracy of Phirbo, BLAST, and WIsH over phage contig length.

**Table S5.** Phage sequence coverage of 485 phages correctly assigned by BLAST and Phirbo to their host species. Sequence coverage was calculated for each phage as the sum of the lengths of its non-overlapping high scoring pairs to the genome of the correct host species, divided by the size of the query-phage genome. Prophages were assumed to have sequence coverage greater than or equal to 50%.

**Table S6.** Summary of taxonomic affiliations of 271 phages that had sequence coverage < 50% with the host species genomes.

**Table S7.** Protein families present in sequence regions of 271 phage genomes that were used by BLAST and/or Phirbo in host prediction. The table provides information on each protein family (prokaryotic Virus Orthologous Group (pVOG)) used by BLAST and Phirbo, including: (i) pVOG description and functional assignment (manually curated), (ii) pVOG count (number of times a given pVOG was present in the phage genome, as well as in sequences used by BLAST or Phirbo), (iii) pVOG percentage (pVOG count divided by pVOG count in the genome), and (iii) *P*-value of pVOG enrichment.

**Table S8**. RNA families present in sequence regions of 271 phage genomes that were used by BLAST and Phirbo in host prediction. The table provides information on each Rfam family used by BLAST and Phirbo.

**Table S9.** F1 scores of Phirbo, VirHostMatcher-Net, and PHP evaluated on the Wang *et al.* data set for 1,462 phages and 62,493 candidate hosts.

**Table S10.** Comparison of Phirbo’s host prediction performance between BLAST-based and Mash-based rankings of prokaryotic species.

## REFERENCES

1. Suttle CA. Marine viruses--major players in the global ecosystem. Nat Rev Microbiol. 2007;5:801–12.

2. Breitbart M, Bonnain C, Malki K, Sawaya NA. Phage puppet masters of the marine microbial realm. Nat Microbiol. 2018;3:754–66.

3. Roux S, Brum JR, Dutilh BE, Sunagawa S, Duhaime MB, Loy A, et al. Ecogenomics and potential biogeochemical impacts of globally abundant ocean viruses. Nature. 2016;537:689–93.

4. Norman JM, Handley SA, Baldridge MT, Droit L, Liu CY, Keller BC, et al. Disease-specific alterations in the enteric virome in inflammatory bowel disease. Cell. 2015;160:447–60.

5. Manrique P, Bolduc B, Walk ST, van der Oost J, de Vos WM, Young MJ. Healthy human gut phageome. Proc Natl Acad Sci U S A. 2016;113:10400–5.

6. Meyer JR. Sticky bacteriophage protect animal cells. Proceedings of the National Academy of Sciences of the United States of America. 2013;110:10475–6.

7. Reardon S. Phage therapy gets revitalized. Nature. 2014;510:15–6.

8. Salmond GPC, Fineran PC. A century of the phage: past, present and future. Nat Rev Microbiol. 2015;13:777–86.

9. Svoboda E. Bacteria-eating viruses could provide a route to stability in cystic fibrosis. Nature. 2020;583:S8–9.

10. Dedrick RM, Guerrero-Bustamante CA, Garlena RA, Russell DA, Ford K, Harris K, et al. Engineered bacteriophages for treatment of a patient with a disseminated drug-resistant Mycobacterium abscessus. Nat Med. 2019;25:730–3.

11. Samson JE, Moineau S. Bacteriophages in food fermentations: new frontiers in a continuous arms race. Annu Rev Food Sci Technol. 2013;4:347–68.

12. Sulakvelidze A. Using lytic bacteriophages to eliminate or significantly reduce contamination of food by foodborne bacterial pathogens. J Sci Food Agric. 2013;93:3137–46.

13. Paez-Espino D, Eloe-Fadrosh EA, Pavlopoulos GA, Thomas AD, Huntemann M, Mikhailova N, et al. Uncovering earth’s virome. Nature. 2016;536:425–30.

14. Edwards RA, McNair K, Faust K, Raes J, Dutilh BE. Computational approaches to predict bacteriophage–host relationships. FEMS Microbiol Rev. 2016;40:258–72.

15. Ahlgren NA, Ren J, Lu YY, Fuhrman JA, Sun F. Alignment-free d_2^* oligonucleotide frequency dissimilarity measure improves prediction of hosts from metagenomically-derived viral sequences. Nucleic Acids Res. 2017;45:39–53.

16. Galiez C, Siebert M, Enault F, Vincent J, Söding J. WIsH: who is the host? Predicting prokaryotic hosts from metagenomic phage contigs. Bioinformatics. 2017;33:3113–4.

17. Andersson AF, Banfield JF. Virus population dynamics and acquired virus resistance in natural microbial communities. Science. 2008;320:1047–50.

18. Wang W, Ren J, Tang K, Dart E, Ignacio-Espinoza JC, Fuhrman JA, et al. A network-based integrated framework for predicting virus-prokaryote interactions. NAR Genom Bioinform. 2020;2:lqaa044.

19. Lu C, Zhang Z, Cai Z, Zhu Z, Qiu Y, Wu A, et al. Prokaryotic virus host predictor: a Gaussian model for host prediction of prokaryotic viruses in metagenomics. BMC Biol. 2021;19:5.

20. Altschul SF, Madden TL, Schäffer AA, Zhang J, Zhang Z, Miller W, et al. Gapped BLAST and PSI-BLAST: a new generation of protein database search programs. Nucleic Acids Res. 1997;25:3389–402.

21. Lima-Mendez G, Faust K, Henry N, Decelle J, Colin S, Carcillo F, et al. Ocean plankton. Determinants of community structure in the global plankton interactome. Science. 2015;348:1262073.

22. Flores CO, Meyer JR, Valverde S, Farr L, Weitz JS. Statistical structure of host-phage interactions. Proc Natl Acad Sci U S A. 2011;108:E288–97.

23. Webber W, Moffat A, Zobel J. A similarity measure for indefinite rankings. ACM Trans Inf Syst. 2010;28:1–38.

24. Saito T, Rehmsmeier M. The precision-recall plot is more informative than the ROC plot when evaluating binary classifiers on imbalanced datasets. PLoS One. 2015;10:e0118432.

25. Davis J, Goadrich M. The relationship between Precision-Recall and ROC curves. In: Proceedings of the 23rd international conference on Machine learning - ICML ’06. New York, New York, USA: ACM Press; 2006. doi:10.1145/1143844.1143874.

26. Villarroel J, Kleinheinz KA, Jurtz VI, Zschach H, Lund O, Nielsen M, et al. HostPhinder: A phage host prediction tool. Viruses. 2016;8. doi:10.3390/v8050116.

27. Ondov BD, Treangen TJ, Melsted P, Mallonee AB, Bergman NH, Koren S, et al. Mash: fast genome and metagenome distance estimation using MinHash. Genome Biol. 2016;17. doi:10.1186/s13059-016-0997-x.

28. Gao NL, Zhang C, Zhang Z, Hu S, Lercher MJ, Zhao X-M, et al. MVP: a microbe–phage interaction database. Nucleic Acids Res. 2018;46:D700–7.

29. Paez-Espino D, Roux S, Chen I-MA, Palaniappan K, Ratner A, Chu K, et al. IMG/VR v.2.0: an integrated data management and analysis system for cultivated and environmental viral genomes. Nucleic Acids Res. 2019;47:D678–86.

30. Roux S, Hallam SJ, Woyke T, Sullivan MB. Viral dark matter and virus-host interactions resolved from publicly available microbial genomes. Elife. 2015;4. doi:10.7554/eLife.08490.

31. Lawrence JG, Ochman H. Amelioration of bacterial genomes: rates of change and exchange. J Mol Evol. 1997;44:383–97.

32. Pride DT, Wassenaar TM, Ghose C, Blaser MJ. Evidence of host-virus co-evolution in tetranucleotide usage patterns of bacteriophages and eukaryotic viruses. BMC Genomics. 2006;7:8.

33. Carbone A. Codon bias is a major factor explaining phage evolution in translationally biased hosts. J Mol Evol. 2008;66:210–23.

34. Sharp PM, Rogers MS, McConnell DJ. Selection pressures on codon usage in the complete genome of bacteriophage T7. J Mol Evol. 1984;21:150–60.

35. Morgado S, Vicente AC. Global in-silico scenario of tRNA genes and their organization in virus genomes. Viruses. 2019;11:180.

36. Sousa JAM de, Pfeifer E, Touchon M, Rocha EPC. Genome diversification via genetic exchanges between temperate and virulent bacteriophages. bioRxiv. 2020. doi:10.1101/2020.04.14.041137.

37. Shapiro JW, Putonti C. Gene co-occurrence networks reflect bacteriophage ecology and evolution. MBio. 2018;9. doi:10.1128/mbio.01870-17.

38. Hernandes Coutinho F, Zaragosa-Solas A, López-Pérez M, Barylski J, Zielezinski A, Dutilh BE, et al. RaFAH: A superior method for virus-host prediction. bioRxiv. 2020. doi:10.1101/2020.09.25.313155.

39. Camacho C, Coulouris G, Avagyan V, Ma N, Papadopoulos J, Bealer K, et al. BLAST+: architecture and applications. BMC Bioinformatics. 2009;10:421.

40. Pedregosa F, Varoquaux G, Gramfort A, Michel V, Thirion B, Grisel O, et al. Scikit-learn: Machine Learning in Python. J Mach Learn Res. 2011;12:2825–30.

41. Grazziotin AL, Koonin EV, Kristensen DM. Prokaryotic Virus Orthologous Groups (pVOGs): a resource for comparative genomics and protein family annotation. Nucleic Acids Res. 2017;45:D491–8.

42. Kalvari I, Nawrocki EP, Ontiveros-Palacios N, Argasinska J, Lamkiewicz K, Marz M, et al. Rfam 14: expanded coverage of metagenomic, viral and microRNA families. Nucleic Acids Res. 2020. doi:10.1093/nar/gkaa1047.

43. Rice P, Longden I, Bleasby A. EMBOSS: The European molecular biology open software suite. Trends Genet. 2000;16:276–7.

44. Finn RD, Clements J, Eddy SR. HMMER web server: interactive sequence similarity searching. Nucleic Acids Res. 2011;39 Web Server issue:W29–37.

45. Nawrocki EP, Eddy SR. Infernal 1.1: 100-fold faster RNA homology searches. Bioinformatics. 2013;29:2933–5.

46. Virtanen P, Gommers R, Oliphant TE, Haberland M, Reddy T, Cournapeau D, et al. SciPy 1.0: fundamental algorithms for scientific computing in Python. Nat Methods. 2020;17:261–72.

