## Additional file 1 for "Taxonomy-aware, sequence similarity ranking reliably predicts phage-host relationships"

### Suplementarny Information for

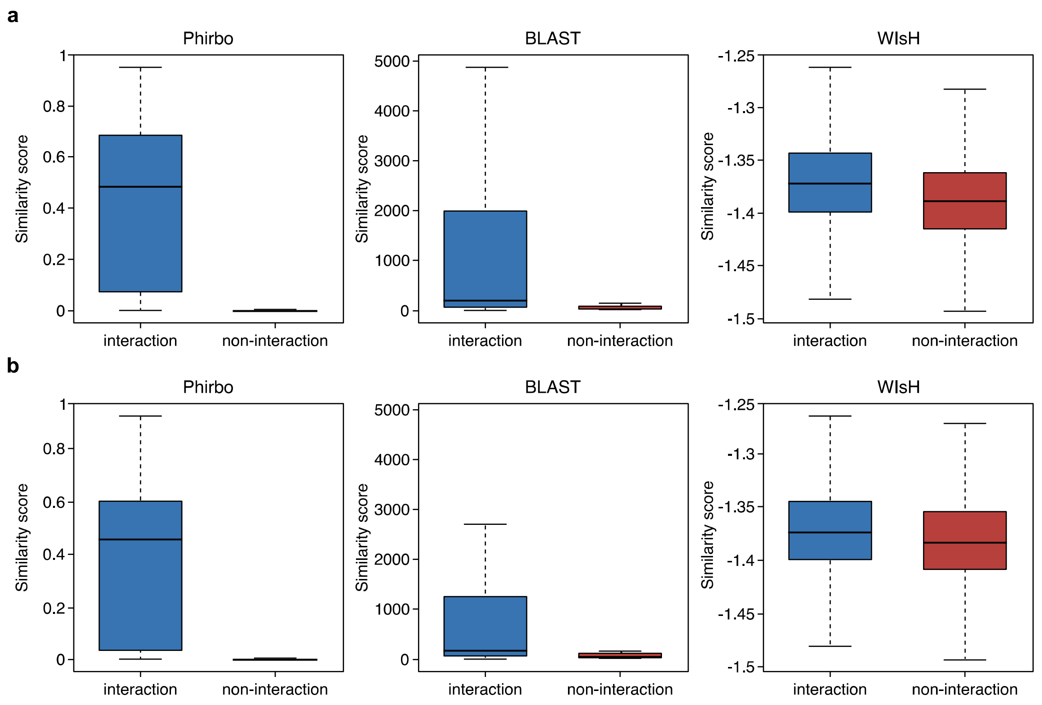


**Figure S1.** Discriminatory power of Phirbo, BLAST, and WIsH scores to differentiate between interacting and non-interacting phage-host pairs. Phage-host pairs were obtained from **a.** Edwards *et al.* and **b.** Galiez *et al.* data sets. Box plots show the distribution of scores for all interacting phage-host pairs ($n$ = 16,757 and $n$ = 26,024 in Edwards *et al.* and Galiez *et al.*, respectively) and the same number of randomly selected, non-interacting phage-host pairs. The horizontal line in each box displays the median; boxes display the first and third quartiles; whiskers depict lowest and highest non-outlier scores (details of distributions including outliers are provided in **Additional file 1: Table S1**).

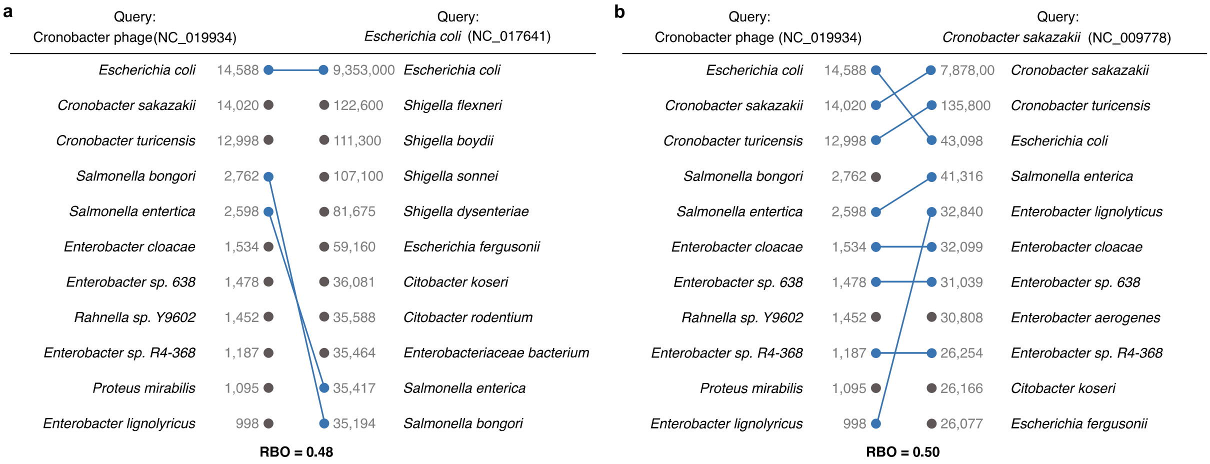


**Figure S2.** Host predictions for Cronobacter phage ENT39118 (RefSeq accession: NC_019934) using **a.** BLAST and **b.** Phirbo. Querying the Cronobacter phage sequence with a BLAST search against the host database returned the genomic sequence of *Escherichia coli* (NC_017641) as the best match (bit-score = 14,588), and *Cronobacter sakazakii* (NC_009778) as the second-best match (bit-score = 14,020). Phirbo predicted *Cronobacter sakazakii* as the top-score host for the Cronobacter phage due to the highest extent of overlap between the top-ranking BLAST matches of each sequence (NC_019934 and NC_009778) of the same database. For clarity, only the first ten BLAST matches are shown.

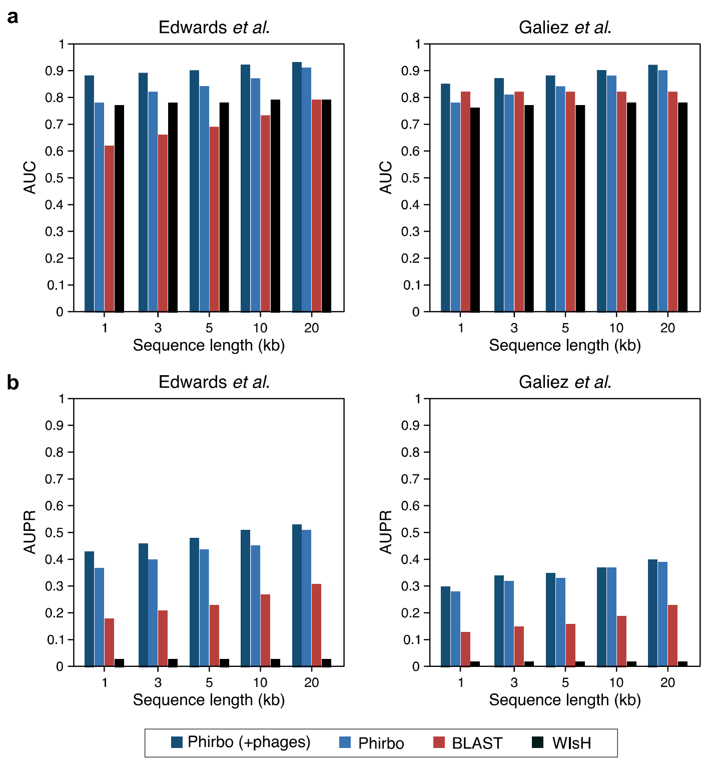


**Figure S3.** Host prediction performance of Phirbo, BLAST and WIsH over phage contig length in terms of **a.** Area under the curve (AUC) and **b.** Area under the precision-recall curve (AUPR). Bars indicate the AUC or AUPR averaged across 10 replicates at a given subsampling length of phage sequence.

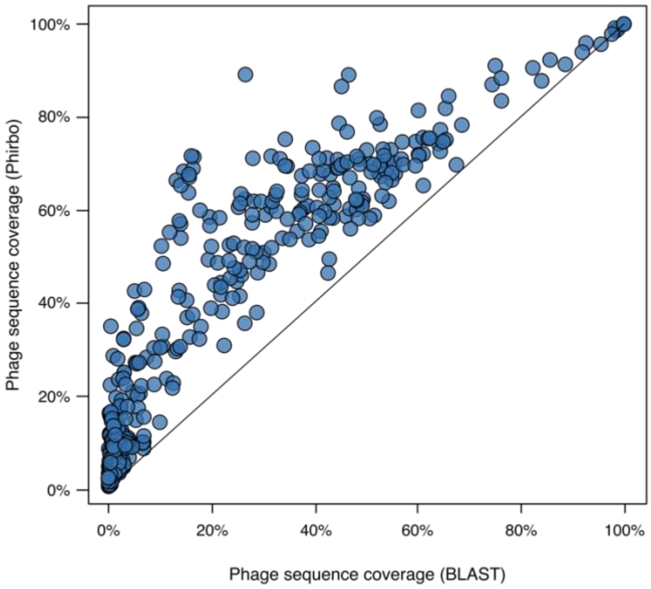


**Figure S4.** Scatter plot of the phage sequence coverage used in host predictions of Phirbo versus that of BLAST. Each dot represents a phage genome.
